# Droplet microfluidic PicoSorter for high throughput and active selection of cellulolytic microorganisms

**DOI:** 10.1101/2025.09.20.677519

**Authors:** Luca Potenza, Łukasz Kozon, Lukasz Drewniak, Tomasz S. Kaminski

## Abstract

Classical enrichment methods for microorganisms rely on growth in selective media, but such practices are relatively expensive, low-throughput, and may result in a biased representation of taxa. Alternatively, microorganisms can be cultivated in thousands of picoliter droplets of equal volume, with the most efficient strains selected through quantitative assays at unprecedented ultrahigh throughput. Here, we present a novel high-throughput microfluidic technology for the characterization of cellulolytic microbial communities at the single-cell level. Individual microbial cells are encapsulated in picoliter droplets for clonal cultivation, after which a colorimetric assay using Congo red is applied to identify positive droplets containing cellulose-degrading strains. These positive droplets are then actively sorted at high throughput using absorbance-activated droplet sorting. The critical component of the enrichment assay we propose is the PicoSorter module, which expands the range of available droplet-based enrichment methods. The platform introduces a new design that enables buffer picoinjection required for the assay, followed by droplet sorting on a millisecond timescale.

**Highlights:**

- Development of a novel device (PicoSorter) capable of performing multiple microfluidic operations simultaneously.
- Implementation of a colorimetric droplet-based assay for the detection of cellulose.
- The proposed absorbance-based HTS enables active and efficient enrichment of cellulolytic bacteria.

## Introduction

### Microbial cellulolytic activity in modern biotechnological applications

Cellulolytic activity is crucial for lignocellulose transformation into biofuel and valuable products like furfural, isoprene, sorbitol, and xylitol^1^. Such types of hydrolytic activity is also used in soil bioremediation, where microorganisms are introduced to degrade pollutants^2^. The microbial cellulolytic activity supports bioremediation processes by aiding in pollution cleanup. Waste lignocellulosic substrates, like sawdust and hay, improve soil permeability and provide enzymes like laccases and peroxidases that break down hydrocarbons^3^. Given the relevance of this type of hydrolytic activity, today there is a high demand for tools to isolate cellulolytic microbial consortia.

### Current methods for characterization of cellulolytic activity and high-throughput screening platforms

Despite the importance of cellulolytic microorganisms for various biotech applications, the selection of highly efficient species from the environment remains challenging. Screening of cellulolytic activity has been executed for decades using Congo red agar plates. The assay is based on a selective medium containing carboxymethylcellulose (CMC) as the primary carbon source, and it is supplemented with the Congo red dye that binds to the biopolymer. Upon bacterial growth, a clear halo surrounding the colonies indicates their cellulolytic activity^4^. Aside from the direct use of Congo red agar plates, various methodologies for quantitative screening of cellulolytic activity comprises mainly colorimetric assays that detect increasing concentrations of the reducing sugars. Standard protocols for the detection of cellulases are based on absorbance measurements - e.g. the 3,5-dinitro salicylic acid (DNS) assays for quantification of reducing sugars concentration^5^. This common assay, however, shows one major limitation - different reducing sugars exhibit variable colorimetric responses. The DNS method can be used as an accurate protocol for the detection of reducing sugar only when glucose is the only monomer constituting the polymer.

However, to date, none of the abovementioned commonly used techniques has been implemented in a high-throughput droplet microfluidic format^6^. Such an approach allows the execution of microbial cultivation and biochemical assays inside stable microdroplets^7,8^, where small volumes, ranging from single pico- to a few microliters, ensure a substantial reduction of time and costs per screening. Currently, it is possible to generate microdroplets on a chip, merge and split them, control their chemical contents, and monitor in situ the outcome of the assay with various detection modes. The feasibility of single-cell encapsulation followed by clonal cultivation is one of the main advantages of this approach, which allows researchers to characterize rare or slow-growing taxa that are very difficult to enrich via classical methods^9–11^.

The first demonstration of a droplet-based protocol for screening cellulolytic bacteria has been implemented via a flow-cytometry workflow. The assay was developed based on fluorescence-activated cell sorting (FACS) and polydisperse bulk double emulsions^12^. Cellulolytic activity is quantified via coupled reactions that generate hydrogen peroxide resulting from the hydrolysis of cellulose. The researchers used a reference library with different percentages of cellulase-active cells and demonstrated a 12-fold enrichment of the positive cellulolytic samples. A different study described the microfluidic enrichment of cellulolytic microbes isolated from wheat stubble. The assay relies on the hydrolysis of a fluorogenic substrate composed of cellobiose linked to a 6,8-difluoro-7-hydroxycoumarin-4-methane-sulfonate^13^. Results demonstrated that the enriched population of microorganisms, isolated by droplets, exhibited 17- and 7-fold higher cellobiohydrolase and endoglucanase activities, respectively. The enriched consortium is significantly different in taxonomic diversity and richness compared to the microbiota isolated with classical protocols. Similarly, a different droplet microfluidic assay for cellulolytic screening has been described based on the formation of a fluorescent product in a coupled reaction that leads to the cleavage of aminophenyl fluorescein (APF) molecules into fluorescein^14^. The assay successfully enriches cellulases-expressing yeasts from a set of reference libraries after a short incubation time of 20 minutes from cell encapsulation.

In this manuscript, we propose the use of absorbance-activated droplet sorting (AADS)^15,16^ to quantify the cellulolytic activity by measuring the cellulose content left after microbial growth, regardless of the type of cellulolytic enzymes that a strain can secrete within a droplet. The optical assay is executed on the newly developed PicoSorter, a microfluidic platform that integrates droplet reinjection, picoinjection, transmittance measurement, and sorting within a single chip, enabling high-throughput operation on a millisecond timescale.

## Materials and methods

### Bacterial strains

Two reference strains were cultivated to validate our PicoSorter: *Escherichia coli* DH5α (the Leibniz Institute DSMZ collection) and *Cellulosimicrobium cellulans* pw7-13 (isolated from the wastewater treatment plant in Poland, Lubuskie voivodeship). Bacterial samples are stored as a collection of strains in a -80°C freezer in 2 mL cryovial tubes. A cryoprotective solution (30% glycerol in 20% DMSO, prepared in ddH_2_O, v/v) is mixed at a 1:1 ratio with an overnight LB culture and then frozen.

### Preparation of the bacterial suspension

Bacterial strains were revitalized from overnight cultures grown on solid LB at 30°C overnight. The LB medium contained 1.5% (w/v) agar plus 20 g LB Broth Difco Miller in 1 L of ddH_2_O. Single colonies were then scraped and inoculated in flasks containing 10 mL of liquid LB medium and incubated overnight under shaking at 150 rpm at 30°C.

The optical density was determined by a plate reader measuring the absorbance values at 600 nm wavelength (OD600). Cultures were diluted appropriately with the liquid BH-CMC 2.5% medium to reach the desired droplet occupancy. The cell loading concentration was adjusted to achieve a Poisson distribution with an average occupancy (λ) of x bacteria per droplet, where λ represents the mean number of bacteria encapsulated per droplet. The BH-CMC was autoclaved for 15 minutes at ¾ atm, and its composition for 1 L preparation is listed as follows: 1 g KH_2_PO4, 1 g K_2_HPO_4_, 1 g (NH4)_2_SO_4_, 0.2 g MgSO_4_, 0.02 g CaCl_2_, 0.05 g FeCl_3_, 10 g peptone, CMC 25 g/l.

### Fabrication of microfluidic devices

The microfluidic devices used in this study were designed in AutoCAD (Autodesk), and master molds were generated through standard photolithography. Polydimethylsiloxane (PDMS) devices were then fabricated via soft lithography. A description of the fabrication process is provided in the Supporting Information.

### Droplet generation and incubation

Flow-focusing droplet generators (50 µm width, 50 µm height) were placed on the stage of an inverted optical microscope (Olympus CKX53). Two 1 mL gas-tight glass syringes and one 2.5 mL gas-tight glass syringe (Hamilton) were filled with BH-CMC 2.5% medium and with Novec HFE-7500 oil (3M) supplemented with 2% RAN fluorosurfactant (RAN Biotechnologies), respectively. All these syringes were firmly locked in dosing modules of a syringe pump (Nemesys low-pressure module, Cetoni) and connected to the inlet of the chip by 10-15 cm PTFE tubing (0.5 mm I.D., 1.0 mm O.D., Bola Bohlender). The syringes were controlled in real time by Qmix Elements software (Cetoni). Generation of 50 pl droplets at a throughput of approximately 1800 droplets per second was executed at a flow rate of oil of 600 µL/h and 320 µL/h for the syringes containing the culture medium. Droplets were collected from chips in storage chambers containing the same oil and surfactant concentration as described above. Droplets entered the chamber from the tip of the Eppendorf tube. Excess oil flowed from the lower outlet of the chamber due to a higher density compared to the aqueous phase of droplets.

### Droplet pico-injection and sorting

The PicoSorter chip was positioned on the movable stage of an inverted microscope. A total of five syringes were placed on the syringe pump rig, the first two 250 µL syringes for the microbial sample in BH CMC medium (CMC 2.5% + Congo red 3.5 mg/mL), and for the MOPS/NaCl buffer (0.1M MOPS pH 7 and 1 M NaCl). The first spacing oil (1% v/v RAN) and the RI-matching oil composed of 40% v/v refractive index–matching compound (1,3-Bis(trifluoromethyl)-5-bromobenzene, Apollo Scientific) in HFE-7500 were delivered to the chip using 2.5 mL syringes. Lastly, a 1 mL syringe of bias oil was connected to the chip (2% v/v of RAN fluorosurfactant in HFE-7500 oil). Flow rates for the PicoSorter were set as follows: sample 120 µL/h, first spacing oil 300 µL/h, buffer 120 µL/h, RI-matching oil 2400 µL/h, and bias oil 1500 µL/h.

PTFE tubings from the syringes were firmly connected to the PicoSorter inlets.

Fabrication of the PicoSorter chip required integrating incident light and detection multimode optical fibers with FC/PC connectors. Both the incident light and detection fibers had a 125 µm cladding diameter, 50 µm core, and 0.22 NA (M14L05, Thorlabs). Fibers were stripped, once the PVC coating and the protective acrylate layer were removed, the fiber tips were cleaved with a ceramic fiber scribe (CSW12-5, Thorlabs)^17^. Cleavage quality was verified with an inspection of the shape of the beam, repeating if necessary until a spherical light beam was observed. Fibers were then fixed to the microfluidic chip at least a day before experiments. Positioning was verified using a microscope camera as fibers were manually inserted into PDMS-filled channels. Additional stabilization was achieved by attaching fibers to the glass slide with epoxy glue (2031-1, Araldite). The chip was left overnight to cure at room temperature.

### Post-sorting droplet imaging

To observe and assess the type of droplets being enriched via the PicoSorter, the tubing from the positive outlet was connected to a 15 mL Falcon tube containing 0.5 mL of 5% RAN in HFE oil. Once a red rim of droplets - visible due to the Congo red in the BH-CMC medium - accumulated inside the tube, a few µL were gently pipetted into a Glasstic slides (Kova) prefilled with 5% RAN oil to prevent droplet merging. Images were then acquired using the AX-100 camera using PFV4 software (Photron) and saved for further analysis.

## Results and Discussion

### Design the PicoSorter module for multistep droplet assays

To detect microbial cellulolytic activity using an optical assay in microdroplet format, we decided to use Congo red dye, which binds to long-chain biopolymers, as cellulose, resulting in its absorbance shift. Similarly to the agar plate method, this assay can be executed in liquid format in a microtiter well plate^18^. Therefore, we decided to scale down the whole protocol in a microdroplet format. The novel microfluidic workflow we propose offers advantages, such as the feasibility of a high-throughput, quantitative assay for cellulose detection in an aqueous picoliter droplet without the necessity of using fluorescent substrates.

Cellulolytic activity is assessed by quantifying cellulose concentration through a well-established colorimetric reaction, which provides a general measure of hydrolytic capacity rather than selectively targeting the activity of a specific cellulase class, as described in other microfluidic enrichment approaches. The absorbance shift of the Congo red-CMC complex is quantified via a spectrophotometric analysis that detects the absorbance shift at 530 nm wavelength^18^. The microfluidic droplet sorter has been developed based on the previous demonstrations of absorbance-activated droplet sorters (AADS)^15,19^. Since the Congo red assay is based on a multiple-step protocol, we expanded and integrated the AADS design with an additional pico-injection module for the facile addition of buffers to droplets. To the best of our knowledge, this is the first demonstration of picoinjection and sorting integrated on the same microfluidic chip, hereafter referred to as PicoSorter. Traditionally, these operations are executed by two separate microfluidic devices^20^, which results in a longer assay duration and greater losses of droplets during transfer between microdevices.

First, single cells are encapsulated inside 50 pl droplets via a flow-focusing device. Then, droplets are collected and incubated off-chip at 25°C inside droplet chambers to allow microbial growth. A constant flow of oil is provided via a peristaltic pump to provide oxygenation of droplets^21^. To prevent air bubbles from accumulating in the chambers, a bubble trap tube is positioned between the pump and the droplets^22,23^. Afterwards, droplets are reinjected in the PicoSorter device, where first picoinjection of MOPS/NaCl buffer is performed, as described in the “Materials and methods” section. Next, the droplet transmittance is measured by optic fibers aligned across the main channel. Transmittance values are measured as voltage signals, which are then compared against user-defined thresholds to trigger the sorting of droplets containing cellulolytic cultures. All 3 operations, picoinjection, transmittance measurement and droplet sorting, are performed sequentially within the same microfluidic device in milliseconds (Video S1). The chip is placed on the stage of an optical inverted microscope, droplets and microchannels are closely monitored using a high-speed camera connected to the PC. In the upper section of the chip, where picoinjection is executed, a constant electric field is applied using a function generator and a high-voltage amplifier. After transmittance detection, the photodetector sends signals to an Arduino microcontroller, which initiates droplet sorting. The voltage interval used to trigger sorting of cellulolytic cultures can be finely adjusted at any moment during the experiment via the code controlling the Arduino board. Additionally, we utilized manual controls (knobs) on the pulse generator to adjust parameters such as delay and width of the electric pulse. These adjustments are essential to ensure the accurate sorting of single positive droplets. Simultaneously, the signal is sent to a data acquisition device (DAQ) connected to a LabVIEW script running on a PC. The program includes a real-time console for continuous monitoring of the sorting process and data integration for further analysis^15,24^.

We optimized various parts of the PicoSorter design (Figure 2A) to accomplish optimal assay performance in terms of throughput and system stability. To embed the optic fibers in the chip, the device was designed with a double-layer layout. The first layer, with a channel depth of 40 µm, supports evenly spaced droplet flow and picoinjection. In this layer, the channel width expands from 35 to 50 µm to accommodate the formation of 100 pL droplets. The second layer, which contains the AADS section, features deeper channels of 90 µm to enable the insertion of optical fibers. The reinjection chamber of the device is surrounded by a moat channel (ground electrode) to shield droplets from the electric field and prevent the unwanted merging of the emulsion coming from the chamber. In addition, a serpentine channel has been introduced after picoinjection to better mix a MOPS/NaCl reaction buffer with the droplets^25^. The channel providing buffer intersects with the droplet channel in front of the first set of electrodes, where a constant electric field is applied. After picoinjection, droplets pass through a mixing channel, and they are further spaced with oil before final transmittance detection. To reduce edge scattering from droplets, thus enhancing transmittance detection, 40% v/v of 1,3-bis(trifluoromethyl)-5-bromobenzene refractive index (RI) matching compound is diluted in perfluorinated HFE-7500 oil, hence we used the name RI-matching oil^19^. In this application of the PicoSorter device, the RI compound is added exclusively to the RI-matching oil flowing from the channels before droplet detection. The transmittance is measured as a voltage signal that triggers the second set of electrodes. The addition of RI compound to the oil might cause emulsion instability and wetting of channels and outlet tubing, as shown in video S3. To prevent such issues in the sorting section and positive outlet, a stream of so-called bias oil is delivered to enhance efficient droplet sorting (Figure 2B). The bias oil at operating rates flows in a laminar manner alongside the oil from the main channel and fills the positive outlet, thereby preventing RI oil from mixing with positively sorted droplets.

**Figure 1.**
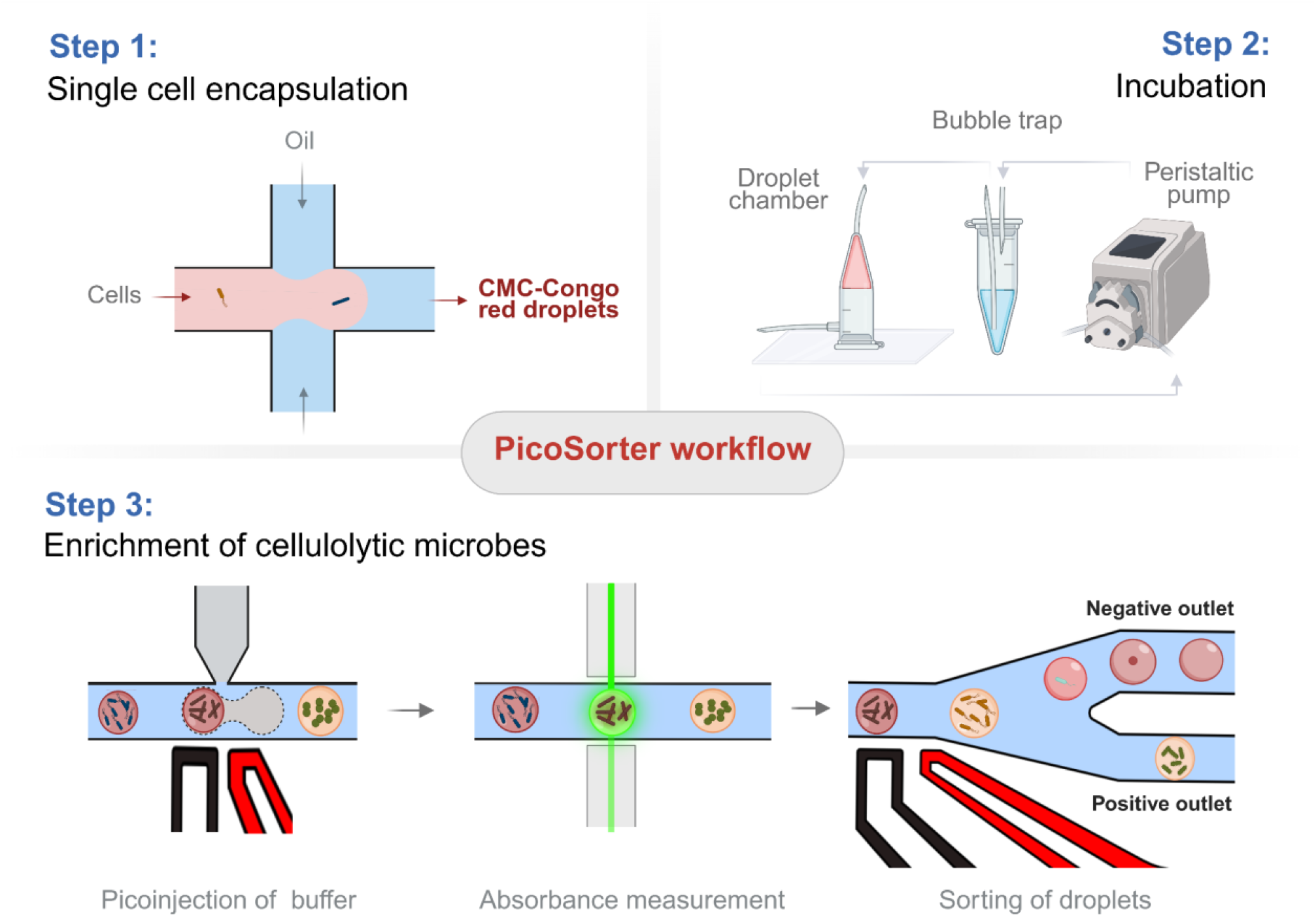
Schematic of the high-throughput screening of cellulolytic activity in microdroplets. The consecutive phases of the microfluidic absorbance-based assay are divided into three steps. Single cells are encapsulated in 50 pl droplets using a flow-focusing device (Step 1), followed by off-chip incubation in chambers with oxygenation via an oil flow regulated by a peristaltic pump (Step 2). Air bubbles are removed with a bubble trap tube. Droplets are then reinjected into a PicoSorter for picoinjection of MOPS/NaCl buffer and transmittance measurement, which triggers sorting of cellulolytic cultures (Step 3).

**Figure 2.**
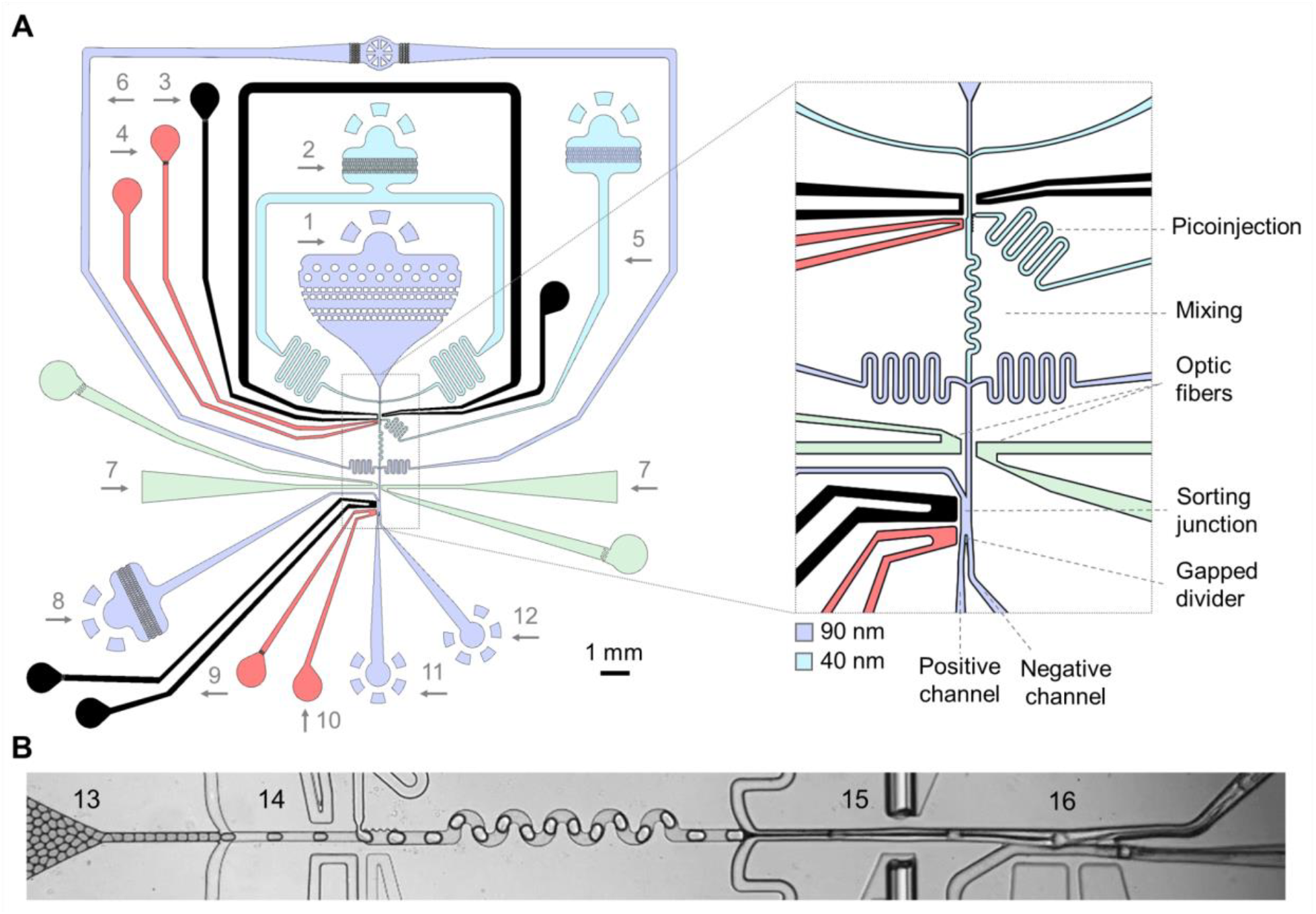
**The design of the PicoSorter double-layered chip for picoinjection combined with absorbance-activated droplet sorting – AADS (A)**. The PicoSorter module includes: 1. Reinjection droplet chamber, 2. Spacing oil channel, 3. Moat channel with ground electrodes for picoinjection, 4. Positive electrode for picoinjection, 5. Picoinjection buffer. 6. RI-matching oil channel, 7. Optical fiber channels, 8. Bias oil channel, 9 Ground electrode for sorting, 10. Positive electrode for sorting, 11. Positive outlet, 12. Negative outlet. Various important features of the PicoSorter device are indicated in the close-up figure. The first layer (light blue) comprises the droplet reinjection and the channel with a height of 40 µm, from where the droplets are brought to the picoinjection section. The second layer (dark blue), with a height of 90 µm, is where the transmittance detection of droplets was executed. **Snapshot of the picoinjection and sorting process (B)**. The microphotograph depicts the consecutive microfluidic steps performed with the PicoSorter: 13. Reinjection of 50 pl droplets, 14. Picoinjection of the buffer required for the assay and mixing, 15. Transmittance detection, 16. Active sorting of 100 pl droplets.

### Demonstration of quantitative measurement of cellulose concentrations

Since Congo red dye is also a well-known pH indicator^26^, it was important to ensure that transmittance measurements from the droplets were not affected by slight changes in pH, thereby guaranteeing reliable detection under microbial cultivation conditions. We first evaluated the influence of pH on the detection of CMC– Congo red in Bushnell Haas (BH) cultivation medium. As shown in Figure 3A, transmittance detection of 0-2.5% CMC remained consistent across the tested pH range. Additional information regarding the colourimetric detection of CMC is provided in the SI section, Figure S3. We also tested the sensitivity of our absorbance-based method using CMC concentrations ranging from 0% up to 2.5% (w/v), as shown in Figure 3B. To enhance transmittance detection in microdroplets, the concentration of Congo red was increased to a maximum of 3.5 mg/mL. This final concentration did not inhibit microbial growth, allowing quantitative cellulose measurements to be performed in picolitre droplets. Picoinjection of 50 pl of buffer (50 mM MOPS, pH 7, 0.5 M NaCl) into each 50 pl droplet was essential for the assay and provided several advantages. Background noise significantly affects the detection of droplets with low CMC content, introducing considerable uncertainty in transmittance measurements. Therefore, picoinjection of droplets with MOPS/NaCl buffer is necessary for accurate detection of low CMC content droplets and/or the presence of cellulolytic cultures. Buffer picoinjection expanded the measurable range of CMC concentrations, and enabled better discrimination between droplets containing cellulolytic strains and negative control droplets, Figure 3 (C). However, owing to the presence of NaCl, the buffer was added only immediately prior to transmittance measurement, as NaCl rapidly promotes Congo red aggregation. This not only interferes with transmittance readings but also tends to clog the filters and inlets, as well as the buffer channel and the picoinjection junction. Overall, the MOPS/NaCl buffer enhanced the precision of transmittance measurements, particularly in droplets containing low CMC concentrations resulting from microbial cellulolytic activity. More detailed results regarding the influence of buffer on the proposed assay can be found in the Supplementary Information, Figure S3.

**Figure 3.**
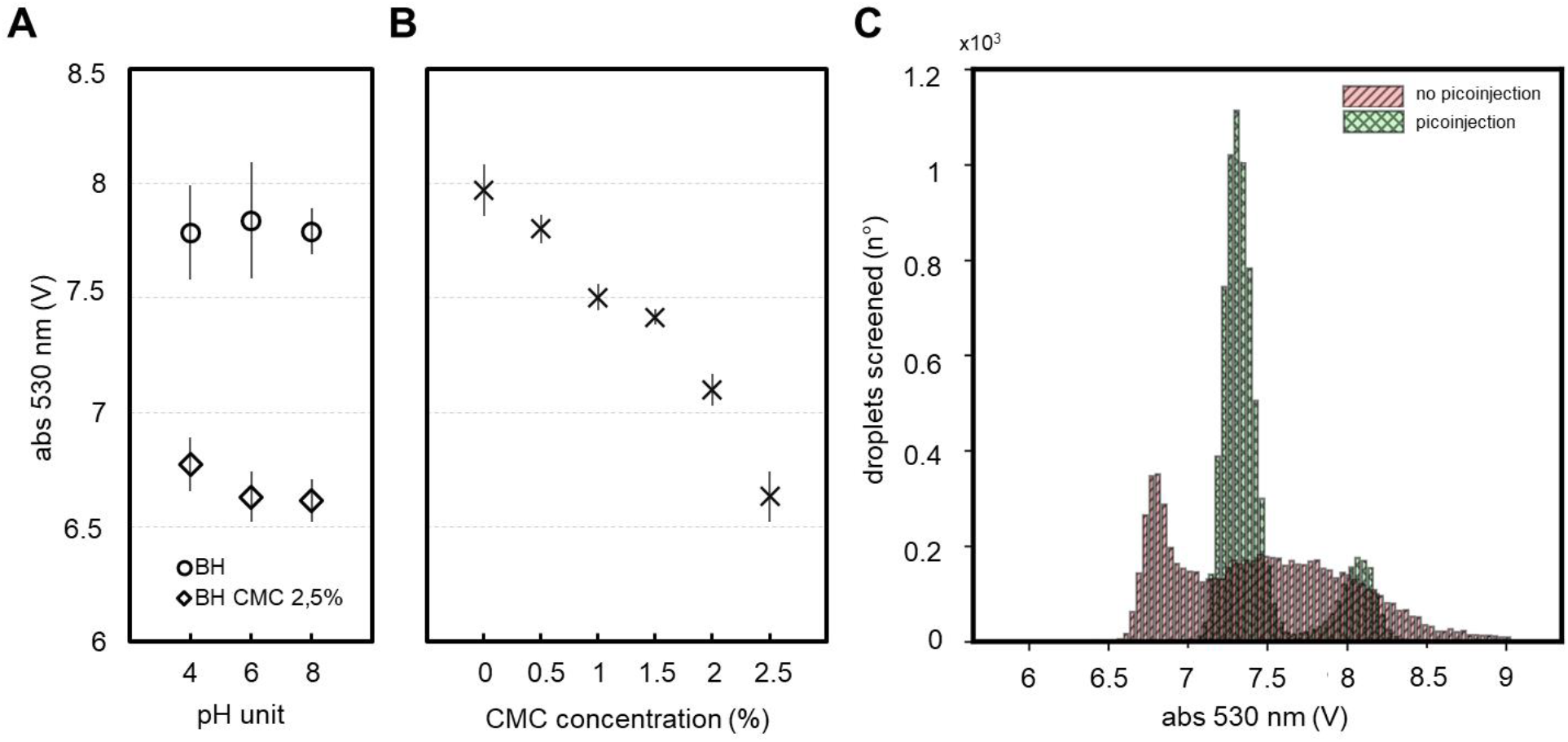
Colorimetric detection of CMC. The scatter plot (**A**) illustrates the relationship between the medium pH and buffer (MOPS/NaCl, pH 7) and the transmittance measurement of CMC content at 530 nm. On-chip detection of 100 pL droplets demonstrated that the microfluidic assay can be effectively performed within the suggested pH range. The scatter plot (**B**) depicts the relationship between CMC concentration and the corresponding absorbance-derived voltage signal at 530 nm, measured at pH 7. An inverse relationship is observed, with increasing CMC concentration resulting in a decrease in the detected voltage signal. The influence of picoinjection of buffer solution on droplet transmittance values is shown in the histogram on the right (**C**), where the absorbance signal of each droplet was quantified and the droplets enumerated. The graphs show results for different droplet populations containing cellulolytic cultures of *C. cellulans* (λ∼0.1). The green histogram represents transmittance measurements of 50 pL droplets that were merged with 50 pL of MOPS/NaCl buffer, while the light red histogram corresponds to 100 pL droplets measured without buffer injection.

### Validation of sorting accuracy by enrichment of cellulolytic *C. cellulans* from a mock microbial consortium

In the final stage of the project, we conducted an enrichment of cellulolytic bacteria cultures derived from single cells that were encapsulated within droplets. The encapsulation of single cells follows a Poisson distribution, resulting in a variable number of cells being present in each droplet while the majority of the droplets remain empty. Two major reference strains were chosen for clonal cultivation in droplets: *E. coli* and *C. cellulans*, which represent, respectively, the non-cellulolytic and the cellulolytic bacteria. These two strains were used to first validate the enrichment of cellulolytic cultures and assess the performance of the PicoSorter as shown in Figure 4.

**Figure 4.**
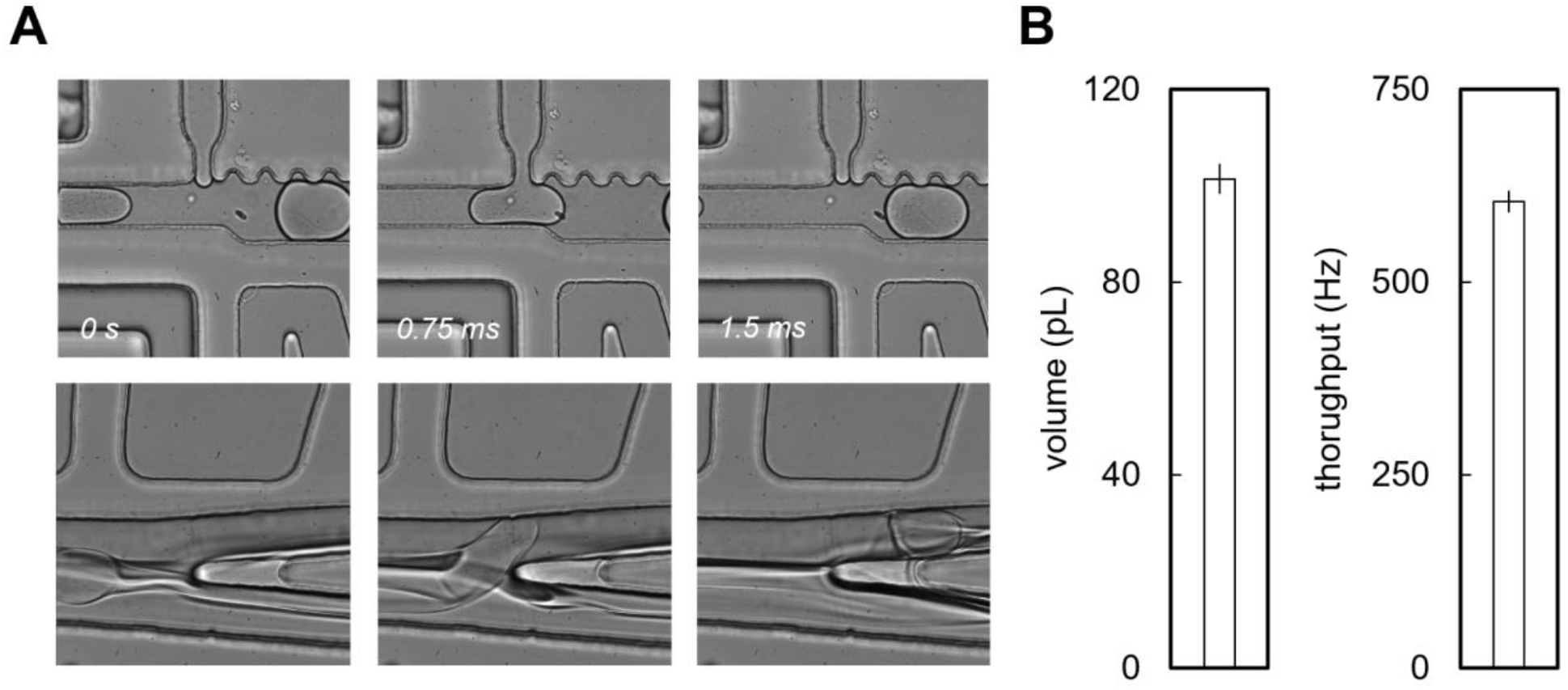
PicoSorter performance under microbial cultivation conditions. The *C. cellulans* emulsion (λ ∼ 0.1) was incubated at 25°C under dynamic droplet conditions and therefore screened. Microfluidic operations simultaneously performed in the PicoSorter device are shown on the left (**A**). A clonal colony of *C. cellulans* growing inside a 50 pL droplet first undergoes picoinjection of buffer, after which the resulting 100 pL droplet is sorted at a throughput of 0.6 kHz (**B**).

To boost microbial growth, we introduced a modified version of the Bushnell Haas (BH) medium supplemented with CMC 2.5% w/v (BH–CMC medium). According to in-bulk cultivation of reference strains, screening of cellulolytic activity can be successfully detected from cultures grown in flasks after 48 hours of incubation at 25°C, as described in Figure S3 of the Supporting Information section. Cultivation of the reference strains under microdroplet conditions required slightly extended incubation. The experimental setup employs dynamic droplet incubation^21^, with a constant oil flow (600 μL/h) supplying oxygen to the droplets of 2% RAN fluorosurfactant in Novec HFE-7500. Under these conditions, microfluidic screening can be successfully performed after 72 hours of incubation at 25 °C. The extended incubation time required for cultivating strains in droplets is likely due to the unique microenvironment within the droplets. Nonetheless, the clonal growth of cells within stable microdroplets enables a more accurate representation of complex microbial communities and proved superior to bulk culturing methods, which are biased towards the fastest-growing microorganisms. After incubation, droplets were reinjected into the PicoSorter to execute absorbance-based CMC detection. Values from empty droplets and non-cellulolytic *E*.*coli* microbes were similar, while the values from *C. cellulans* were generally a unit higher, correlating with the reduced amount of CMC left in the droplets, therefore highlighting the cellulolytic activity of the selected strain, Figure 5 (A, C). The PicoSorter electronics were set to send a pulse and sort positive droplets to the positive outlet if the measured transmittance was within a customizable voltage interval. On the contrary, all the *E*.*coli* and empty droplets were discarded via the negative outlet of the chip.

**Figure 5.**
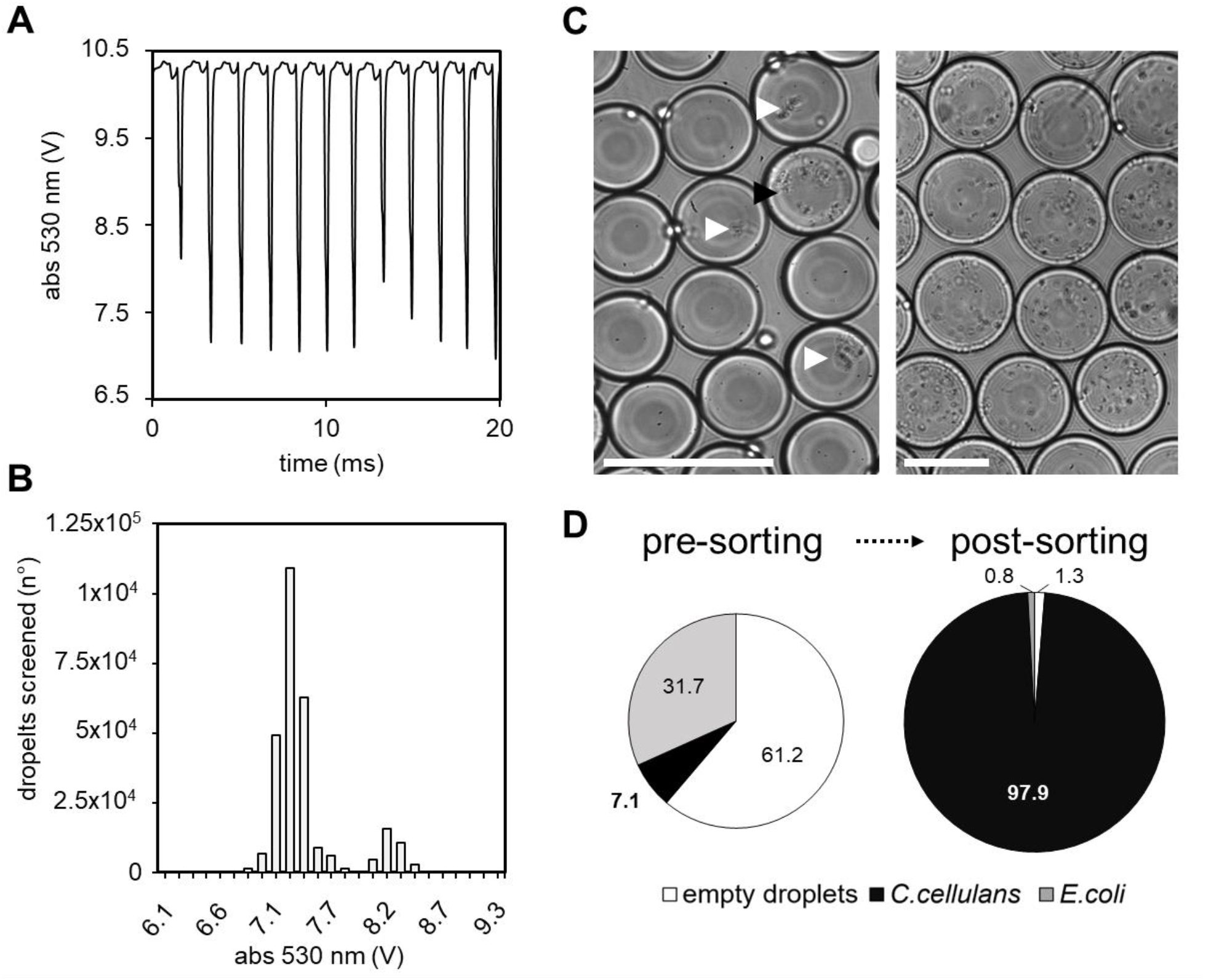
Screening and enrichment of cellulolytic bacteria from a mock microbial consortium. The proposed colorimetric assay for HTS of microbial cellulolytic activity was validated using a binary synthetic community composed by *C*.*cellulans* (λ∼0.1) and *E*.*coli* (λ∼0.3). Droplet voltage signals are displayed in the line plot (**A**), with each peak representing a single droplet. In the histogram, detected peaks are enumerated based on real-time voltage measurements **(B)**. The *C. cellulans* signal (∼8 V) is approximately 1 V higher than droplets containing *E. coli* or empty droplets. Microphotographs on the right (**C**) show the emulsion composition before and after sorting, scale bar 50 µm. The reference strains differ in colony morphology: *E. coli* is indicated by white arrows, while *C. cellulans* droplet is marked with a black arrow. Pre- and post-sorting emulsion compositions (%) were quantified via image analysis and are shown in the pie charts (**D**).

The enrichment factors reported in Table S1 were determined using previously described approaches by Baret et al.^27^ and Zinchenko et al.^28^. The calculated values were approximately 610 and 14, indicating that our approach effectively enriches cellulolytic microorganisms.

## Conclusions

Droplet microfluidics brings several competitive advantages for performing biochemical assays inside stable microdroplets^7^, mainly reduced volumes and costs that allow for the screening of up to 10^8^ samples per day. Droplet technology allows applications like single-cell studies, enabling researchers to screen microcultures originating from a single cell in liquid droplets, making it possible to screen for rare and slow-growing taxa^9^. Here, we proved how the microfluidic system we developed can successfully execute the enrichment of cellulolytic microorganisms based on an absorbance-based assay performed in the PicoSorter device - a novel platform for executing sequential picoinjection and droplet sorting on the same device.

The protocol we propose takes advantage of the high-throughput nature of the droplet microfluidics technique, enabling the sorting of up to 600 droplets per second. Although AADS-based microfluidic protocols have recently achieved sorting rates of up to 2.6 kHz for 50 pL droplets^16^, most current methods do not exceed 0.1–1 kHz^14,15^. Our sorter stands out for its speed, as the PicoSorter combines two distinct functions, setting it apart from most droplet microfluidic assays. Notably, Kaba et al.^16^ reported even higher throughput, reaching 5.4 kHz, but only when operating with ultra-small 10 pL droplets. Such small volumes, however, hinder the growth of bacterial colonies, making them unsuitable for functional screening of microbial activity. In contrast to confocal AADS setups, which require constant objective focus on the sorting region and therefore limit simultaneous control of processes such as picoinjection, the measured values in a fibre-optic detection system are independent of chip positioning.. This flexibility allows the operator to freely monitor and control additional microfluidic operations without compromising sorting performance. Moreover, fibre-based AADS is more cost-effective, does not require advanced microscopy, and is easier to assemble than confocal-based systems. In addition to its current implementation with AADS, the Picosorter system could be adapted to operate with alternative detection modalities. For instance, integration with fluorescence-based readouts^27,29^ (FADS) or scattered light activated droplet sorting^30^ would enable high-sensitivity screening while maintaining compatibility with both label-based and label-free strategies. These potential extensions underscore the versatility of the platform and its applicability across a broad spectrum of droplet microfluidic assays. The PicoSorter could find extensive implementations for various colorimetric reactions that do not require a long incubation time. Independently from the proposed application of this optical assay, the PicoSorter can be implemented for the detection of fast reactions with substrates that leak from droplets to the oil and/or between droplets^31^- i.e., reactions in which the initial rate of product formation is faster than the leakage rate.

Lignocellulosic biomass is an abundant organic resource and is considered a source of renewable energy,^2,32^ however, our knowledge about microbial composition throughout processes like biofuel production and bioremediation is still restricted. There is, in fact, a need for tools to characterize and describe microbial consortia composition, physiology and interactions in an accurate way. The PicoSorter module, combined with the optical assay we described, could find application in screening cellulolytic microbial consortia with a great level of accuracy and performance. Moreover, we foresee the coupling of our new high-throughput screening with a droplet deposition system^33^ to take advantage of the in-droplet cultivation with the upscaling on well plates of each sorted individual microculture^34^. From a protein engineering standpoint, the PicoSorter device could be implemented toward the direct evolution of enzymes to achieve improved hydrolytic activity of cellulases. It is worth mentioning that the colorimetric approach we used does not rely on expensive modules for fluorescence detection, thus expanding the implementation prospects of the PicoSorter module.

Despite the high throughput and sensitivity level of the proposed protocol, the limitation of our method relies on the use of CMC as a substrate instead of pure cellulose. Cellobiohydrolases display low activity on CMC, and for measuring the activity of this class of enzymes, different assays should be considered. During the development of the PicoSorter we tested various designs, including different optics and droplet channel layout. We aimed to enhance the transmittance detection of droplets by introducing an S-shaped channel in front of the optical fibers. This configuration squeezed droplets in front of the optical fibers, aiming to enhance colorimetric detection. Nonetheless, this approach did achieve a limited ability to better distinguish between signals from positive and negative samples. In the future, to better characterize cellulolytic activity in droplets, the dynamic range of absorbance detection can be improved, or the UV-ViS spectrum measured for better resolution of the activity^35^.

The PicoSorter addresses the issues associated with traditional microbiological isolation protocols, opening new possibilities for academic research and biotechnological applications. This method marks significant progress in the field of microbial screening and enrichment, and the device we developed can be applied in different fields of modern environmental microbiology, such as biofuel production, bioremediation, and sustainable industrial processes.

## Supplementary information

The Supporting Information provides detailed protocols for photolithography, fabrication of microfluidic devices, moulds, droplet incubation chambers, the emulsion oxygenation system, and sorting parameters. It also includes procedures for the colorimetric detection of soluble CMC and bulk screening of microbial cellulolytic activity. Supplementary videos illustrate the complete microfluidic assay covering droplet picoinjection, mixing, transmittance measurement, and sorting. The Arduino scripts necessary for the operation and control of the PicoSorter system are publicly accessible via the following repository: https://github.com/Microfluidic-UW/PicoSorter-.git.

- Microfluidic chip design of the droplet generator and the PicoSorter device (DXF).
- Fabrication protocols.
- Video S1. Full PicoSorter view - sorting of cellulolytic *C. cellulans* (MP4).
- Video S2. Screening of *C*.*cellulans* - Flow of droplets in the PicoSorter positive outlet (MP4).
- Video S3. Screening of *C*.*cellulans* - Flow of droplets in the PicoSorter negative outlet (MP4).
- Video S4. Screening of a mock community - close-up of picoinjection (MP4).
- Video S5. Screening of a mock community - close-up of sorting *C*.*cellulans* (MP4).

## Acknowledgements

This research was funded by the TEAM-NET programme of the Foundation for Polish Science no. POIR.04.04.00-00-14E6/18-00 as a part of Measure 4.4 of the 2014–2020 Smart Growth Operational Programme, EU, and by National Science Centre, Poland (grant SONATA BIS no. 2023/50/E/ST4/00545). Research infrastructure used in the project was co-funded by the “*Excellence Initiative – Research University (2020-2026)*” programme via Action I.4.2 “*Fund for the Renovation and Development of Research Infrastructure*”. We thank Kumar Pranaw and Namrata Joshi for providing the cellulolytic strain used to validate the PicoSorter device. The scheme of the PicoSorter workflow, Figure 1, was prepared with BioRender.com.

---

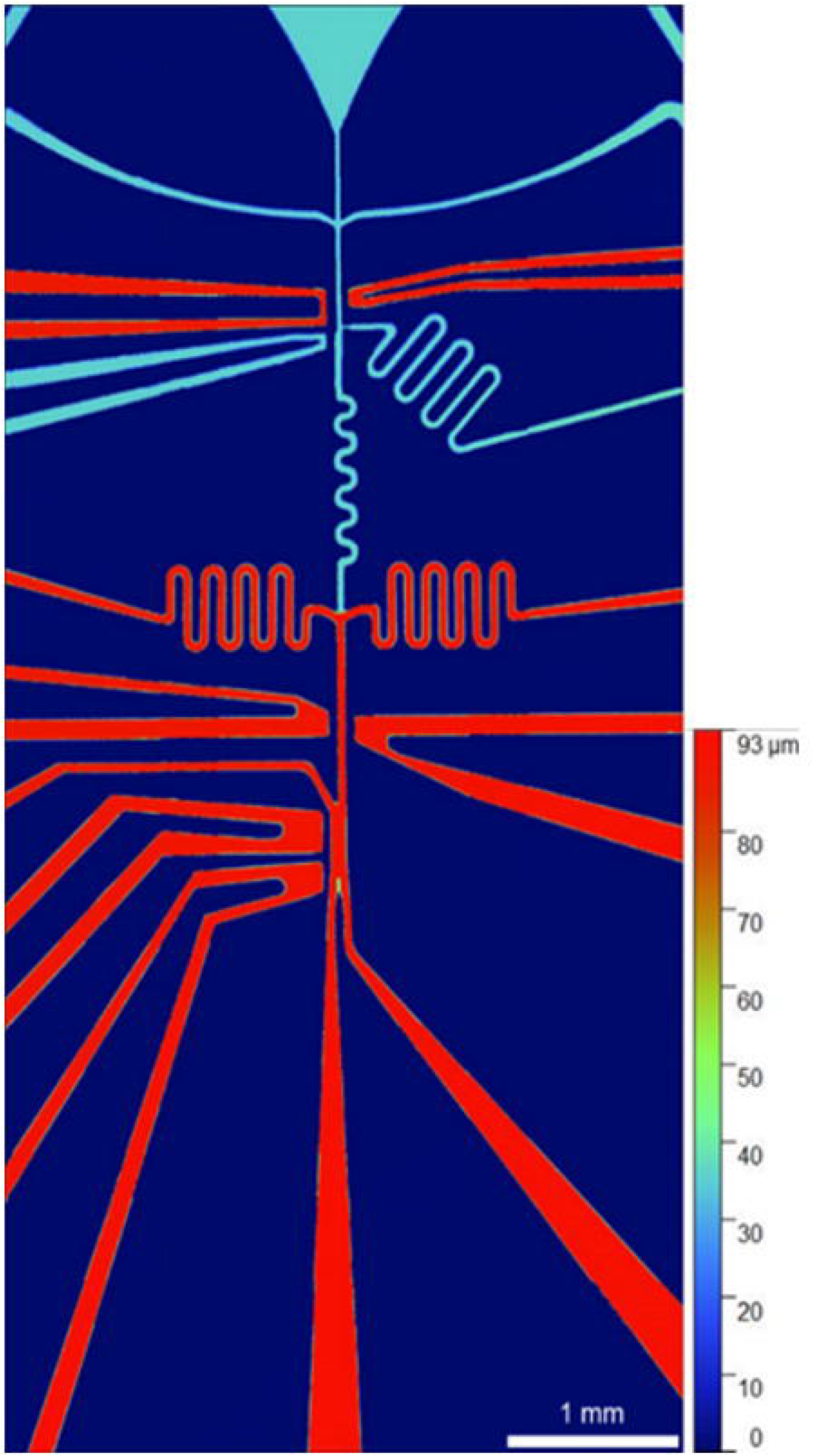

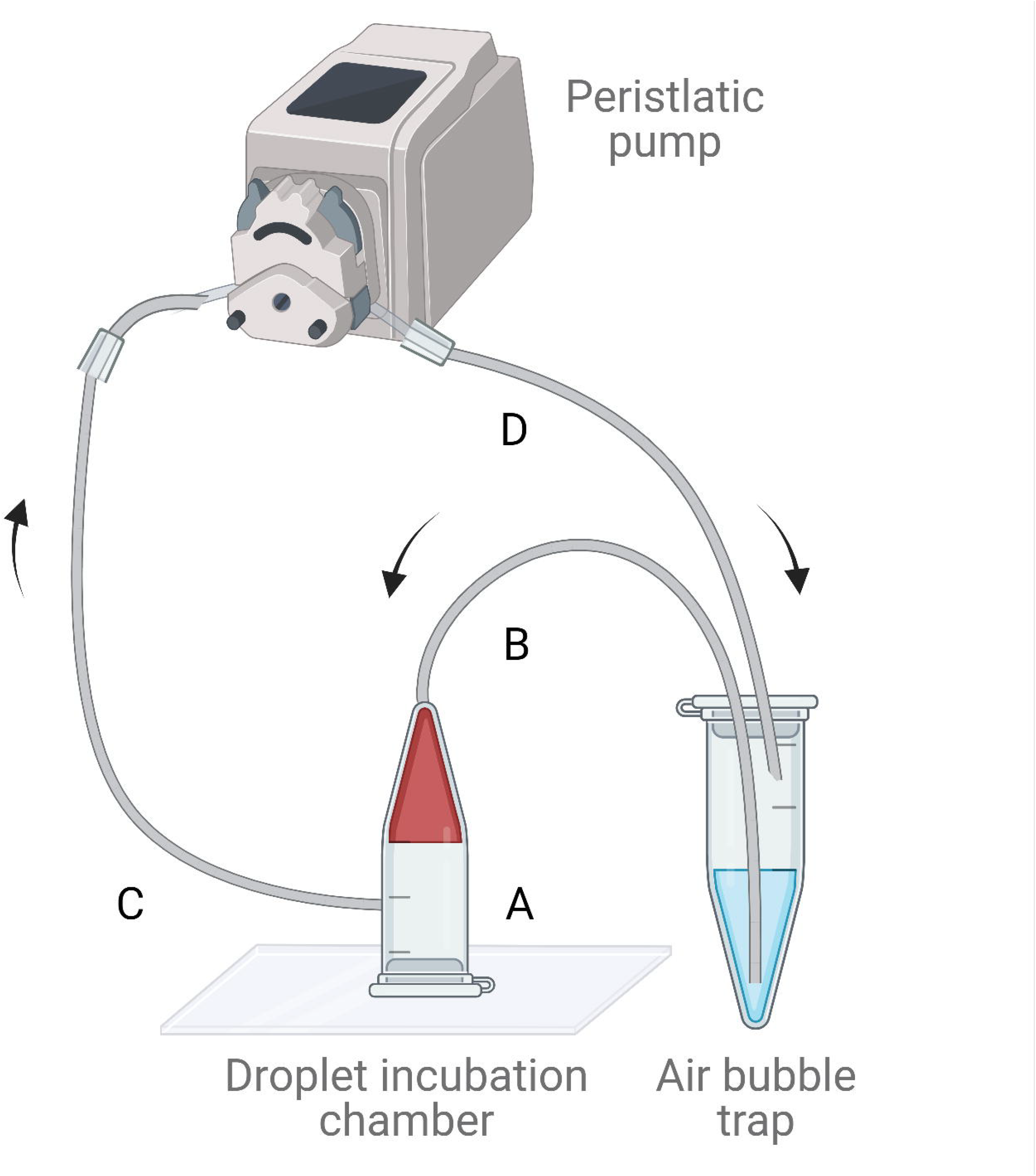

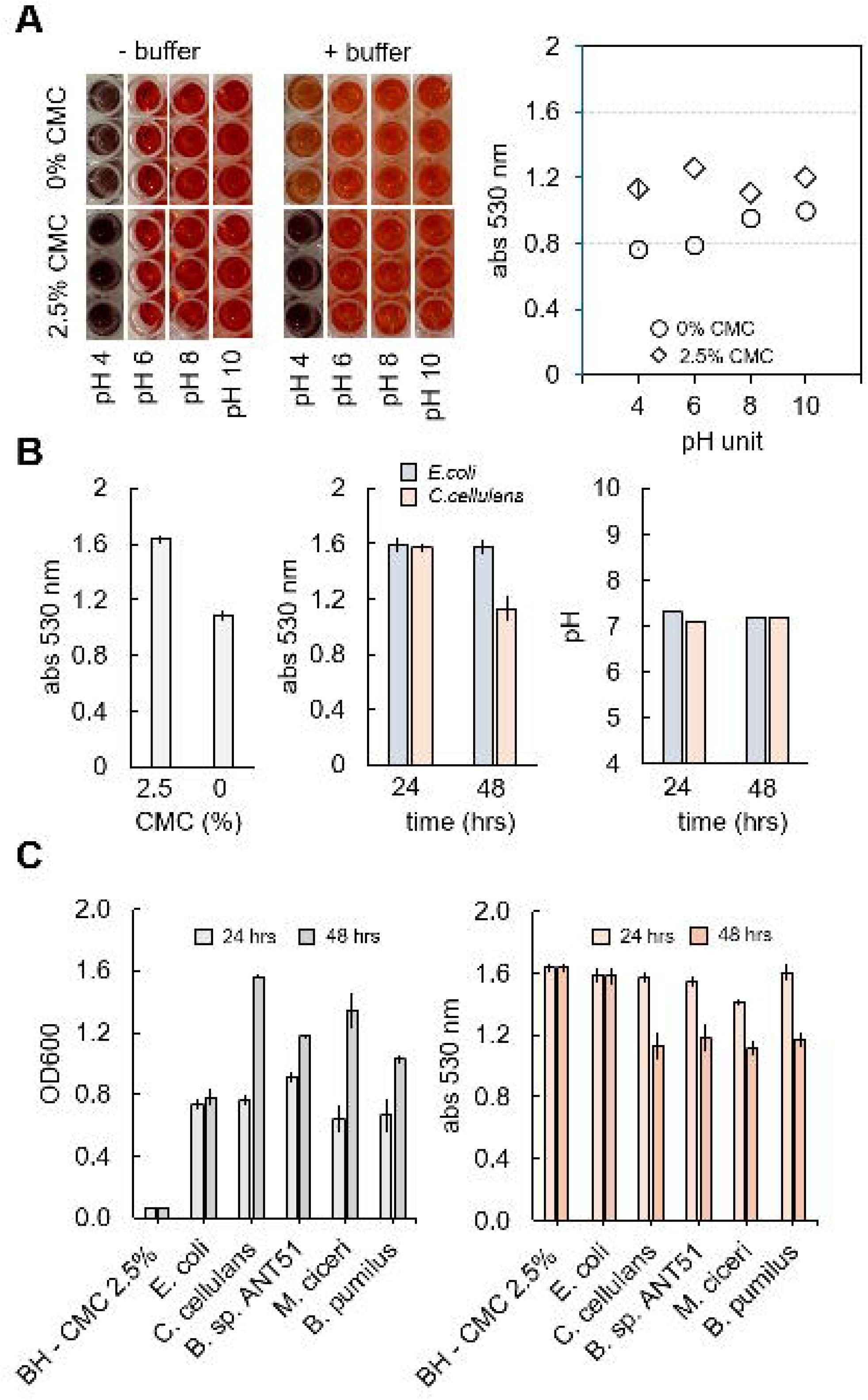

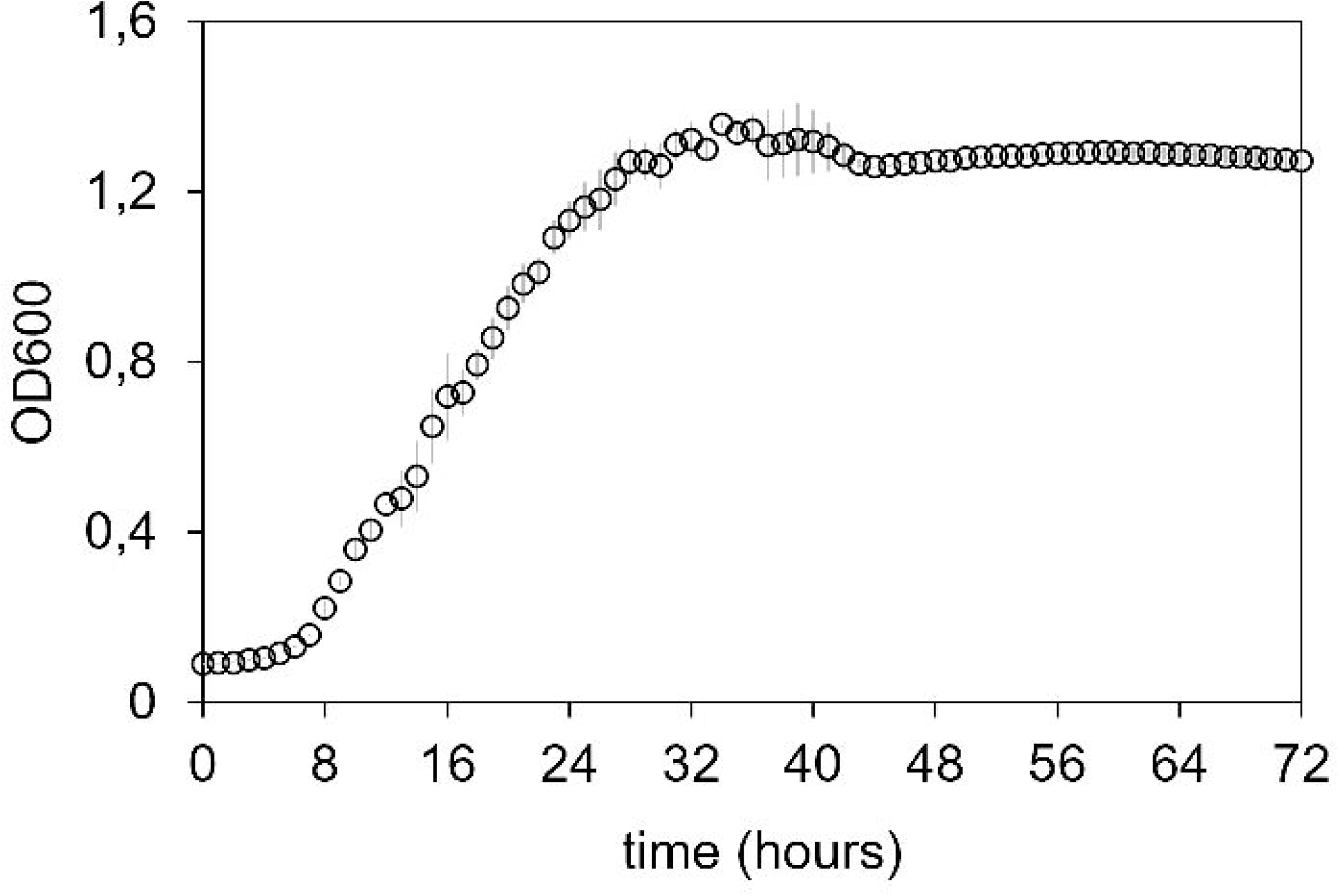

