## Supplementary information for "Droplet microfluidic PicoSorter for high throughput and active selection of cellulolytic microorganisms"

Luca Potenza[a], Łukasz Kozon[a], Lukasz Drewniak[b], Tomasz S. Kaminski[a].
a - Department of Molecular Biology, Institute of Biochemistry, Faculty of Biology, University of Warsaw, Poland.
b - Institute of Bioengineering, Faculty of Biology, University of Warsaw, Poland.

### *Table of Content*

**Supplementary methods**

i. Fabrication of microfluidic devices S2

ii. Photolithography S2

iii. Soft lithography S2

iv. Fabrication of storage chambers for droplet incubation S3

v. Droplet oxygenation during incubation in storage chambers S3

vi. Picoinjection and sorting settings. S5

**Supplementary figures**

i. Figure S1. PicoSorter layout. S2

ii. Figure S2. Oxygenation system layout. S3

iii. Figure S3. Influence of pH on the colorimetric detection of CMC and screening of microbial cellulolytic activity. S4

iv. Figure S4. Growth curve of *C. cellulans* S5

**Fabrication of microfluidic devices.** The microfluidic devices used in this work were designed in AutoCAD (Autodesk), and the master molds were fabricated by standard photolithography and soft lithography[1]. To fabricate PDMS devices, the following steps were executed: first, a master mold containing a designed structure was generated via photolithography, and then the replica containing the channels was produced by pouring PDMS onto the master mold. Replicates were bonded on a glass slide, and the entire chips were silanized by flushing silane to prevent any contact between aqueous droplets and the inner wall of microfluidic channels.

**Photolithography.** Microfluidic molds were created on 3-inches silicon wafers (Microchemicals) using high-resolution acetate masks (Microlithography Services) and SU-8 2050 photoresists patterning (Kayaku Advanced Materials). The SU-8 spin-coated wafers were then exposed to the UV using an MJB4 mask aligner (SÜSS MicroTec)[2]. The CAD designs of the droplet generation device and PicoSorter module are attached in a supplementary .dxf file. The thickness of the structures (corresponding to the depth of channels in the final microfluidic devices) was measured using a GT-Contour profilometer and 5x objective (Bruker). The profile of the droplet sorting module is presented in Figure S1.

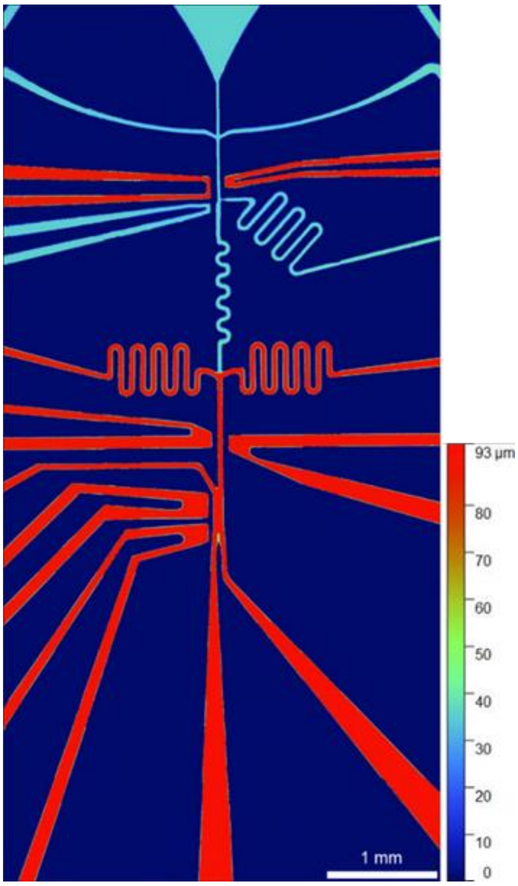


**Figure S1. PicoSorter layout.** Profilometry scan displays a topographical representation of the module surface, highlighting its two-layer structure.

**Soft lithography.** To generate a single microfluidic device, approximately 5 grams of PDMS (Sylgard 184, Dow) are weighed in a plastic cup. The curing agent was added and mixed at a 10:1 (w/w) ratio, and the liquid PDMS was then degassed in a vacuum chamber. PDMS is poured on the SU-8 master wafer, placed in a Petri dish, and cured in the oven at 70°C for 4 hours. Once the PDMS is solidified, the device is peeled off from the mold, and the holes for inlets and outlets are punched using a 1 mm biopsy puncher (Kai Medical). The flow-focusing droplet generators and the Picosorter chips were bound to glass slides (VWR) using an automated plasma system (Zepto, Diener Electronics). The PicoSorter required an additional thin PDMS layer between the glass and the chip, which was used for fibre insertion. First, the thin layer was bounded to the glass, and afterwards the PicoSorter was bounded to the PDMS layer. Hydrophobic modification of the chips was carried out by flushing the devices with a 0.5% v/v solution of trichloro (1H,1H,2H,2H perfluorooctyl)-silane (Sigma-Aldrich) in Novec HFE-7500 oil (3M) and baked on a hot plate at 80°C for 30 minutes to evaporate fluorocarbon liquid.

**Fabrication of storage chambers for droplet incubation.** The chambers were made according to the protocol published by Neun et al.[3] A biopsy punch (Kai Medical) was used to make 1-mm holes on the bottom tip and the side of a 0.5 ml Eppendorf tube. The tube lid was then glued to a 1-mm thick glass slide with cyanoacrylate glue (PR 1500, 3M). Two pieces of 20-cm-long Teflon (PFTE) tubing (0.4mm I.D., 0.9mm O.D., Bola Bohlender) were then inserted into the holes and glued to the surface of an Eppendorf tube. During the droplet generation, the flow-focusing chip was connected to the upper tubing, allowing droplets to be collected in the upper section of the tube, due to their natural buoyancy compared to perfluorinated oil. After incubation, to sort the droplets, the direction of flow was reversed, and the oil was pumped through the side tubing, moving the droplets towards the PicoSorter module via the top tubing.

**Droplet oxygenation during incubation in storage chambers.** Emulsions were incubated for 72 hours by dynamic droplet incubation (DDI), which enhances microbial growth and activity[4]. The setup used by us in this study is presented in Figure S2. Measures were taken to remove any bubbles present in the tubing. The schematic representation of the setup employed for oxygenating the droplets in the experimental workflow. It highlights the crucial role of the peristaltic pump and bubble trap that simultaneously allowed for oxygen delivery and prevented air bubble formation in the incubation chamber.

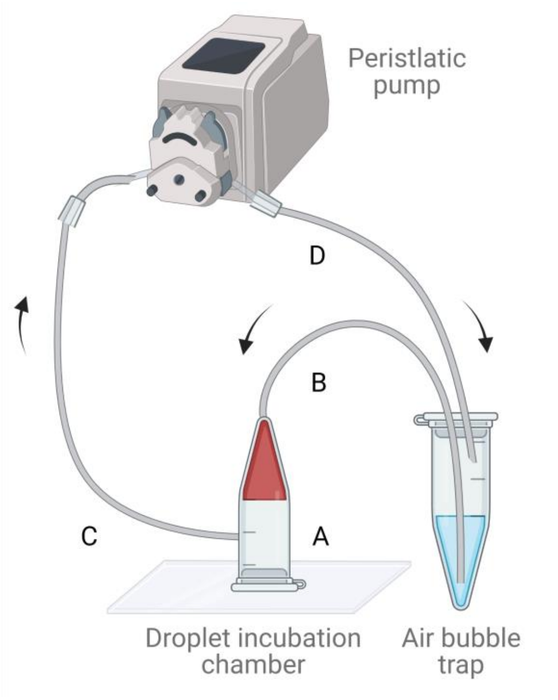


**Figure S2. Oxygenation system layout.** The first 0.5 ml Eppendorf tube was glued to a glass slide and used as a droplet chamber (**A**). The first tubing was glued to the top part of the chamber (**B**). A second tubing was inserted closer to the lid (**C**), Tygon tubing were securely connected via a piece of PTFE tubing to a peristaltic pump (Reglo ICC, Ismatec) using adhesive glue. The other end of the tubing coming from the pump was glued on the lid of a second Eppendorf tube filled with 2% RAN fluorosurfactant solution in filtered Novec HFE-7500 (**D**). This arrangement ensured a continuous flow of the oil phase in a closed system. Oil flow is directed from the top of the tube, where droplets accumulate, to the lower inlet of the chamber at 600 µl/hour. As a result, the emulsion was held in place by buoyancy and prevented from leaking out of the chamber.

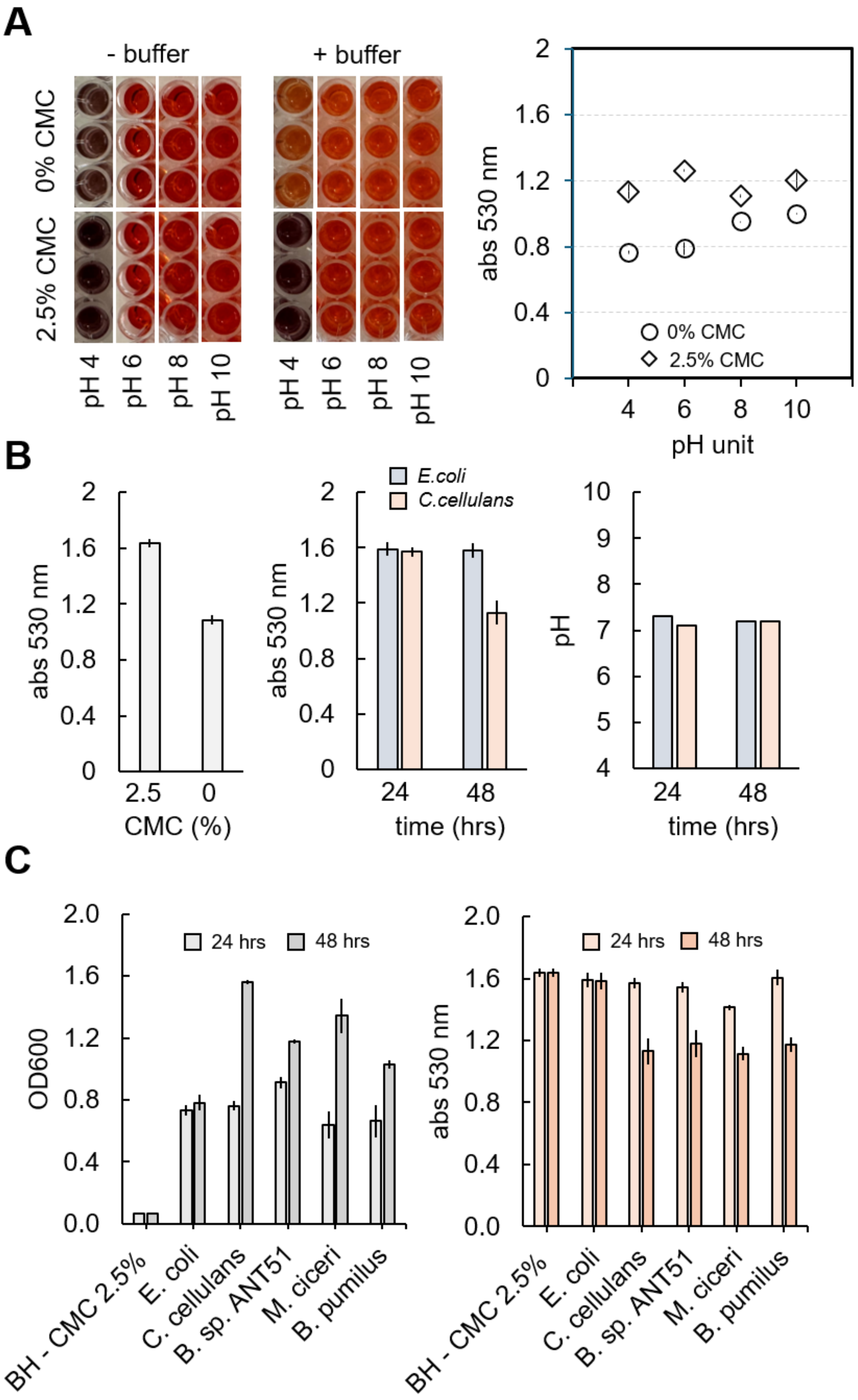


**Figure S3. Influence of pH on the colorimetric detection of CMC and screening of microbial cellulolytic activity.** The influence of pH on the detection of CMC–Congo red was tested across different pH levels in 0–2.5% CMC BH medium (**A**). Below pH 6, Congo red showed aggregation, a phenomenon partially mitigated by the addition of MOPS/NaCl buffer. The scatter plot on the right, obtained from medium supplemented with buffer, indicates that absorbance at 530 nm enables quantitative discrimination between the two CMC concentrations across all tested pH values, although the absorbance range progressively decreases with increasing pH. Complementary bulk assays were conducted to further validate the reliability of the proposed absorbance-based method (**B**). The medium (0–2.5% CMC) was tested as a control, alongside two reference strains: *E. coli* (non-cellulolytic) and *C. cellulans* (cellulolytic). The influence of bulk growth and its correlation with pH was also examined, showing no significant pH variation during bacterial growth and CMC degradation, with values remaining close to the initial medium pH of 7. The column bar plots (**C**) depict the dynamic changes in microbial growth and cellulolytic activity across different reference strains. The experiment included the previously characterised *E. coli* and *C. cellulans*, along with three additional cellulolytic strains: *Bacillus sp. ANT_51*[5] (isolated in Antarctica, from the collection of Institute of Microbiology, University of Warsaw), *Mesorhizobium ciceri*, and *Bacillus pumilus*. Microbial growth is shown in grey bars and expressed as OD600 units, whereas cellulolytic activity is shown in orange and quantified by measuring the residual CMC concentration in the medium. The Congo red assay was performed following the protocol of Haft et al.[6]. Prior to the colorimetric assay, samples were diluted 1:1 with dd$H_2O$ to prevent absorbance values from exceeding 2 units, as higher readings are associated with greater uncertainty.

**Picoinjection and sorting settings.** Picoinjection is carried out by using a first set of a function generator (TGF4042, Aim TTi Ltd) and a high-voltage amplifier (2210, TREK). A square signal is generated and amplified to maintain a constant electric field of 2 volts.
During the sorting process, the voltage signal from the photodetector was recorded with a custom LabVIEW program via a DAQ data acquisition device (Analog-to-Digital Converter USB-6003 Multifunction I/O Device, NI), and at the same time, the same signal was split and sampled in 12-bit resolution via an analog-in pin of an Arduino board microcontroller (Arduino DUE, Arduino). To match the voltage of the detector (10 V) to the maximum voltage sustainable by the microcontroller board (3.3 V), a 10 kΩ resistor was used. In the microcontroller, inputs from the photodetector are compared with a threshold interval. When the signal falls within this range, another pin outputs a 5-volt pulse, triggering droplet sorting. The pulse pace is set to be smaller compared to the droplet interspacing period. Pulse generator (TGP110, Aim TTi Ltd) triggered function generator (TG2000, Aim TTi Ltd ) that executes a square signal in external gated mode, which is next amplified 100 times by a voltage amplifier (2210, TREK) to perform sorting of droplets. Manual adjustments, such as delay and pulse width can be set manually on the pulse generator to achieve optimal sorting of individual positive droplets. Droplet generation and pico-sorting were recorded using a high-speed camera (AX-100, Photron) controlled via the Photron FASTCAM Viewer software (PFV4, Photron).

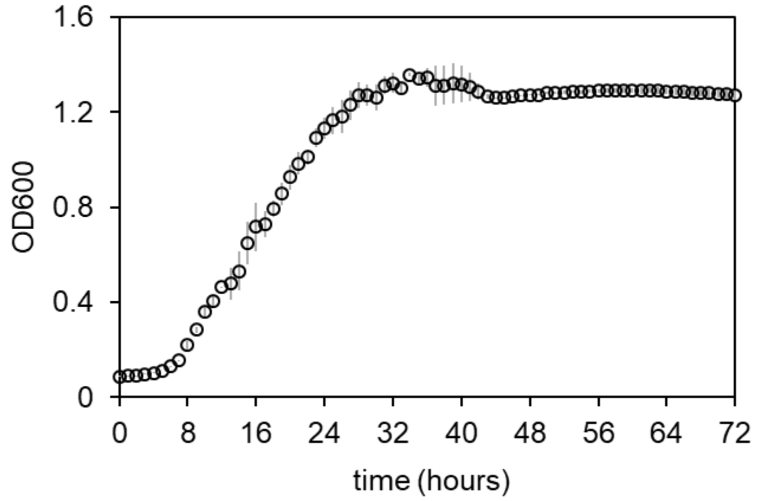


**Figure S4. Growth curve of *C. cellulans*.** The graph shows the growth dynamics of the cellulolytic reference strain cultivated in LB medium in a multiwell plate. The y-axis represents the optical density (OD600) of the bacterial population, while the x-axis represents time (hours). The growth profile exhibits distinct phases: lag (0–8 h), exponential (8–28 h), and a prolonged stationary phase (>28 h). The strain was inoculated into 10 mL of fresh LB medium from an overnight culture (OD600~1) at a 1:1000 (v/v) ratio, transferred to microwells, and cultivated at 25°C under low-speed orbital shaking.

**Determination of enrichment factor.** The enrichment factors described in **Table S1**, were determined using previously described methods by Baret et al.[7] and Zinchenko et al.[8]. The Baret method (**a**) calculates enrichment (**η**) by comparing the final ratio of positive to negative colonies (ε1) after sorting with the initial ratio (ε0) before sorting. The Zinchenko method (**b**) calculates the enrichment factor (**η'**) by comparing the percentage of positive colonies to the total number (ε1') with the initial percentage (ε0').

$$(a) \quad \eta = \frac{\left(\frac{N_+^1}{N_-^1}\right)}{\left(\frac{N_+^0}{N_-^0}\right)} \qquad\qquad (b) \quad \eta' = \frac{\left(\frac{N_+^1}{N_-^1 + N_+^1}\right)}{\left(\frac{N_+^0}{N_-^0 + N_+^0}\right)}$$

The enrichment values we calculated were approximately 610 and 14, indicating that our approach effectively enriches proteolytic microorganisms.

**Table S1. Enrichment values of the PicoSorter protocol.** Droplet classification was performed by visual inspection in Kova chambers before and after sorting.

| Droplets before sorting (n = 652) | | Droplets post sorting (n = 475) | |
|---|---|---|---|
| **% negative** (*E. coli* droplets n = 207,empty droplets n = 399) | **% Positive** (*C.cellulans* droplets n = 46) | **% negative** (*E. coli* n = 4, empty droplets n = 6) | **% Positive** (*C.cellulans* n = 465) |
| **92.9** | **7.1** | **2.1** | **97.9** |

| $N_+^0$ | $N_-^0$ | $N_+^1$ | $N_-^1$ | $\eta$ | $\eta'$ |
|---|---|---|---|---|---|
| 7.1 | 92.9 | 97.9 | 2.1 | **610** | **14** |

### *Videos*

**Video S1. Full PicoSorter view - sorting of cellulolytic *C. cellulans.*** The video shows the active sorting of an emulsion containing *C. cellulans* microcultures originating from single-cell encapsulation (λ~0.1). The oil flow is directed from left to right. Sorting is achieved by applying a transient electric field that diverts positive droplets into the lower outlet - positive channel, while all other droplets (sterile BH–CMC medium) exit through the upper outlet - negative channel. The video was recorded at 8,500 FPS with a 1/20,000 s shutter speed, and the sorting frequency was 0.6 kHz. It was slowed down approximately 350 times.

**Video S2. Screening of *C.cellulans* - Flow of droplets in the PicoSorter positive outlet.** The video shows the sorting of positive droplets in an emulsion containing *C. cellulans* (λ~0.1 positive droplets), focusing exclusively on the positive outlet. The video was recorded at 40× magnification, 10000 FPS, and a 1/100000 s shutter speed. The playback speed was reduced by roughly 2500-fold.

**Video S3. Screening of *C.cellulans* - Flow of droplets in the PicoSorter negative outlet.** The video shows the unsorted fraction of droplets in an emulsion containing *C. cellulans* (λ~0.1 positive droplets), focusing exclusively on the negative outlet. It can be observed that droplets were not stable in RI oil.

The video was recorded at 40× magnification, 10000 FPS, and a 1/100000 s shutter speed. Playback was decelerated by a factor of approximately 100.

**Video S4. Screening of a mock community - close-up of picoinjection.** Picoinjection is shown for an emulsion containing *E. coli* (λ~0.3) and *C. cellulans* (λ~0.1) in 50 pL droplets reinjected from the incubation chamber. A constant electric field induces merging of these droplets with 50 pL of MOPS/NaCl buffer delivered through the side channel. The video was recorded at 10000 FPS with a 1/100000 s shutter speed. The recording was slowed down by a factor of ~750 for visualisation.

**Video S5. Screening of a mock community - close-up of sorting *C.cellulans.*** Sorting is shown for an emulsion containing *E. coli* (λ~0.3) and *C. cellulans* (λ~0.1) in 100 pL droplets obtained after buffer picoinjection and mixing. The droplets flowed in front of the optical fibres, where their absorbance was converted into a voltage signal. Signals within the predefined threshold interval (8.2–8.4 V) triggered a transient electric field that pulled the corresponding droplets into the positive outlet of the PicoSorter. The video was recorded at 10000 FPS with a 1/100000 s shutter speed. It was slowed down approximately 2000 times.
